# An open-source platform for reference data-driven analysis of untargeted metabolomics

**DOI:** 10.1101/2025.11.05.686710

**Authors:** Alejandro Mendoza Cantu, Julia M. Gauglitz, Wout Bittremieux

## Abstract

Untargeted tandem mass spectrometry (MS/MS)-based metabolomics enables broad characterization of small molecules in complex samples, yet the majority of spectra in a typical experiment remain unannotated, limiting biological interpretation. Reference data-driven (RDD) metabolomics addresses this gap by contextualizing spectra through comparison to curated, metadata-annotated reference datasets, allowing inference of spectrum origins without requiring exact structural identification. Here, we present an open-source RDD metabolomics platform comprising a user-friendly web application and a Python software package that perform RDD analyses directly from molecular networking outputs generated by GNPS. The tools support visualization and statistical analysis of RDD results, including interactive bar plots, heat maps, principal component analysis, and Sankey diagrams. We illustrate the approach using a hierarchical reference dataset of 3,500 food items to derive dietary patterns from stool metabolomics data of omnivore and vegan participants. The analysis reveals clear dietary group separation, demonstrating how RDD metabolomics can extract biologically meaningful patterns from otherwise unannotated spectra. Thus, the RDD metabolomics platform removes technical barriers for the metabolomics community to adopting reference data-driven analysis, with the functionality freely available at https://github.com/bittremieuxlab/gnps-rdd and https://gnps-rdd.streamlit.app/.

## Introduction

Untargeted tandem mass spectrometry (MS/MS)-based metabolomics is a widely used technique across the life sciences for comprehensive profiling of small molecules in complex biological systems.^1^ However, despite its power, the complexity of the resulting data and the vastness of the molecular search space mean that only a small fraction of spectra in a typical experiment can be confidently annotated using standard identification approaches such as spectral library matching.^2^ Consequently, a large amount of potentially valuable molecular information remains unexplored.

To help address this limitation, the reference data-driven (RDD) metabolomics approach was recently developed.^3^ Instead of focusing on identifying exact molecular structures, RDD metabolomics contextualizes experimental spectra by comparing them against curated reference datasets. This strategy allows researchers to infer the likely origin or source of a spectrum, even when its precise chemical identity remains unknown.

The value of this approach was illustrated using a large-scale food reference dataset containing more than 3,500 food items analyzed by untargeted metabolomics.^3^ When clinical metabolomics data were compared against this food reference resource, many otherwise unannotated spectra could be linked to dietary origins. RDD metabolomics analysis increased MS/MS spectral usage rates from 5–6% to 15–30%, representing a 3-to 5-fold improvement in data explainability. This, in turn, enabled the reconstruction of dietary patterns directly from plasma and stool metabolomics data.

Although powerful, the initial RDD implementation involved multiple manual programming steps, which posed a barrier to broader adoption by the metabolomics community. To overcome this, we developed an accessible web application that enables researchers to perform RDD analyses directly from molecular networking outputs generated by GNPS^4^ and GNPS2. For users seeking deeper customization, we also provide a dedicated Python software package that extends RDD capabilities to advanced and large-scale analyses. Together, these resources make RDD metabolomics more accessible and reproducible, broadening its potential applications across diverse research domains.

## Methods

### Code availability

The RDD metabolomics Python package is available as open source under the permissive Apache 2.0 license at https://github.com/bittremieuxlab/gnps-rdd and can be installed directly from PyPI using pip. Detailed installation instructions, usage examples, and full application programming interface (API) documentation are provided on the project website. The package relies on NumPy,^5^ Pandas,^6^ Scikit-learn,^7^ and Scikit-bio^8^ for scientific computing, and uses Matplotlib,^9^ Seaborn,^10^ and Plotly^11^ for data visualization. It has been developed following software engineering best practices, including automatic code style enforcement and comprehensive unit testing, with continuous integration to ensure reproducibility and maintainability.

In addition to the Python package, the RDD metabolomics web application is available at https://gnps-rdd.streamlit.app/, and open source on GitHub at https://github.com/AlejandroMC28/gnps_rdd_app/. This application, implemented using Streamlit, provides a graphical interface to the full functionality of the Python package, enabling users to conduct RDD analyses without writing or executing code.

### Software functionality

RDD analyses begin by generating molecular networks on GNPS^4^ or GNPS2, where experimental data are assigned to one group and reference data to another. Step-by-step instructions for setting up this workflow are available in the GNPS documentation (https://ccms-ucsd.github.io/GNPSDocumentation/tutorials/rdd/). Next, the RDD metabolomics software takes as input: (i) the corresponding molecular networking outputs from GNPS or GNPS2, and (ii) a hierarchical metadata sheet linking the reference spectra to their known origins.

Once these inputs are provided, the software interprets the molecular networking results by matching hierarchical metadata to experimental spectra, thereby inferring the origin of the experimental data. The results are exported as an RDD count table, which is a long-format table that links each experimental MS run to specific metadata categories across multiple hierarchical levels, together with the number of spectra assigned to each category. Users can then explore these metadata assignments through a variety of visualization options, including bar plots, box plots, and heat maps, which summarize spectral counts across categories.

The package and web app also support multivariate analysis, such as principal component analysis (PCA), with the option to apply a centered log-ratio transformation for compositional data. Additionally, a Sankey diagram generator is provided to visualize the flow of spectral assignments through the hierarchical metadata system. All plots are interactive, and the display can be customized to adjust the level of granularity and subset of data shown. The plotting backend can be switched between Matplotlib and Plotly with a single line of code or a button in the application, enabling both static, publication-ready visualizations and dynamic, interactive figures.

## Results

To demonstrate the capabilities of the RDD metabolomics web application and Python package, we reproduced selected analyses from the original RDD study,^3^ illustrating how food reference data can be leveraged to derive dietary readouts from clinical metabolomics measurements (**Figure 1**). The reference dataset comprised 3,500 food items spanning diverse food types and sample categories, organized into a hierarchical ontology (e.g., fruit → citrus → lemon → pink lemon). This hierarchical structure enables queries at varying levels of specificity, from broad food classes to individual items. The food reference data and metadata are publicly available on MassIVE (dataset identifier MSV000084900).

**Figure 1.**
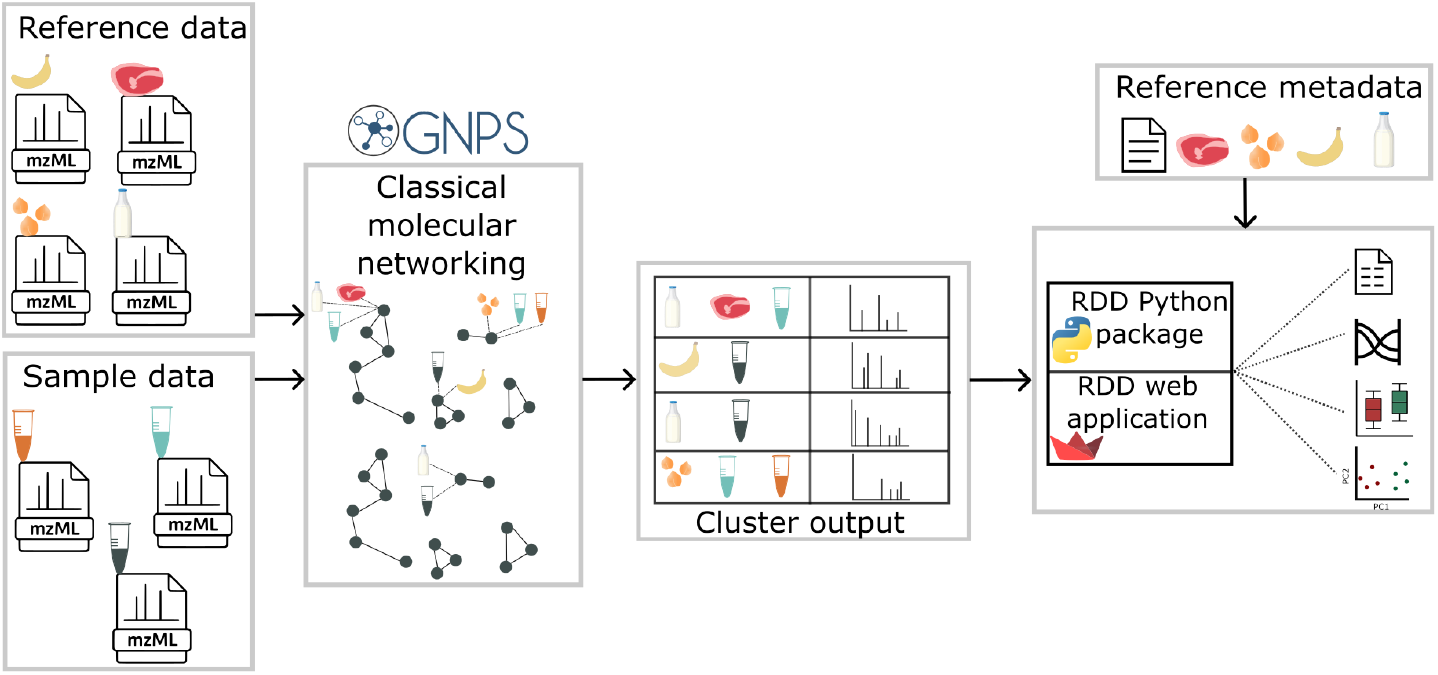
Overview of the RDD analysis workflow. An RDD analysis starts from ontology-controlled source reference data and experimental MS/MS data, which are jointly analyzed using classical molecular networking on the GNPS or GNPS2 platform. Molecular networking output results can be directly imported into the RDD web application to produce an RDD count table, which can further be visualized and analyzed within the web interface. Alternatively, the RDD Python package provides similar functionality for custom data analyses.

For the clinical dataset, we analyzed stool samples from 18 omnivore and 18 vegan study participants (MassIVE dataset identifier MSV00008698). Experimental data were jointly molecularly networked with the food reference dataset on GNPS (GNPS task ID: 74089e95b8df41b2af7c289869dc866f). The resulting molecular networking outputs were then imported directly into the RDD metabolomics web app, which provides an immediate framework for interpreting the spectra in terms of their dietary origins.

Analysis of the RDD count tables revealed expected dietary differences between the two participant groups, with higher proportions of meat, seafood, and dairy assignments in omnivores, and increased fruit and vegetable assignments in vegans. These comparisons can be interactively visualized as box plots or bar plots, with flexible selection of ontology levels and food categories (**Figure 2a**). To capture broader patterns, we applied PCA to the full food count profiles, which produced a clear separation between omnivore and vegan participants based on their RDD-derived dietary signatures (**Figure 2b**). Additionally, a Sankey diagram provided a visual overview of how spectral assignments propagate through the hierarchical food ontology, offering an intuitive depiction of category relationships (**Figure 2c**).

**Figure 2.**
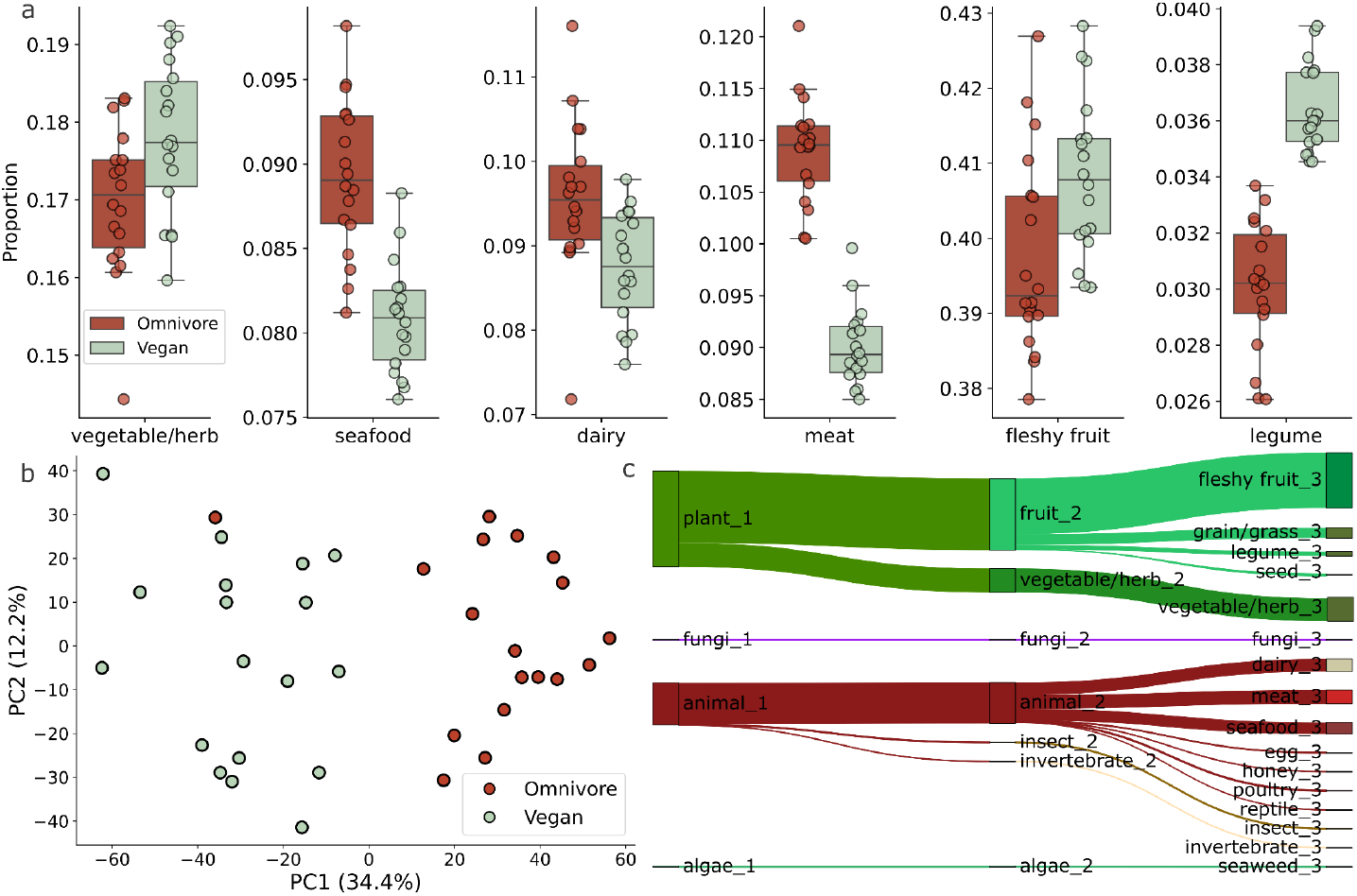
Obtaining dietary readouts from clinical metabolomics data using RDD metabolomics. **(a)** Box plots showing the proportion of spectra assigned to selected food categories in stool samples from omnivore (n = 18) and vegan (n = 18) participants. Food assignments were derived by comparing experimental spectra to a reference dataset of 3,500 foods organized in a hierarchical ontology. Categories shown include meat, seafood, dairy, fruit, legumes, and vegetables, all from the level 3 ontology. Boxes represent the interquartile range, whiskers indicate 1.5 times the interquartile range, and points show individual participants. **(b)** PCA plot based on centered log-ratio transformed RDD count tables for omnivore and vegan participants. Each point represents a participant’s dietary profile inferred from RDD analysis. The separation along the first principal component reflects differences in the consumption of animal-derived versus plant-based foods. **(c)** Sankey diagram illustrating the flow of spectral assignments from high-level ontology categories (e.g., animal product, plant product) to more specific subcategories (e.g., meat → cow, fleshy fruit → berry). The link thickness reflects the proportion of assignments passed from one level to the next, while the diagram is truncated at level 3 of the ontology for visualization purposes.

All of these analyses can be conducted entirely within the web application without the need for programming, while the Python package enables equivalent workflows, often in a single line of code, along with advanced data processing options for more specialized use cases. For example, all analyses presented here can be easily recreated using a Jupyter notebook available on GitHub at https://github.com/bittremieuxlab/gnps-rdd.

## Conclusion

We have developed the RDD metabolomics web application and Python package to enable reference data-driven analyses of untargeted metabolomics datasets. Rather than focusing on identifying exact molecular structures, RDD metabolomics interprets experimental spectra by comparing them to metadata-annotated reference datasets, thereby contextualizing the data in terms of their likely origins.

We illustrated the utility of this approach by deriving dietary readouts from a clinical metabolomics study, demonstrating a clear separation between omnivore and vegan participants. These readouts can be produced directly within the RDD metabolomics tools, as the food reference dataset is included by default, allowing rapid application to dietary studies without additional data preparation.

Although the current implementation comes with preloaded food metadata, the software can incorporate any appropriately formatted reference dataset. Detailed documentation is provided for preparing and integrating custom metadata, which opens the door to a wide range of applications. Potential extensions include mapping the human exposome through reference datasets of personal care products, pharmaceuticals, and environmental chemicals; obtaining microbiome composition readouts from environmental samples; or many other applications. By combining flexible data input with an intuitive interface and robust computational capabilities, the RDD metabolomics platform makes these advanced contextualization analyses broadly accessible to the metabolomics community.

## Acknowledgements

This work was supported by the Research Foundation – Flanders (FWO G0AGQ24N).

## References

1. Aksenov, A. A., Da Silva, R., Knight, R., Lopes, N. P. & Dorrestein, P. C. Global chemical analysis of biology by mass spectrometry. Nat. Rev. Chem. 1, 0054 (2017).

2. Bittremieux, W. et al. Open access repository-scale propagated nearest neighbor suspect spectral library for untargeted metabolomics. Nat. Commun. 14, 8488 (2023).

3. Gauglitz, J. M. et al. Enhancing untargeted metabolomics using metadata-based source annotation. Nat. Biotechnol. 40, 1774–1779 (2022).

4. Wang, M. et al. Sharing and community curation of mass spectrometry data with Global Natural Products Social Molecular Networking. Nat. Biotechnol. 34, 828–837 (2016).

5. Harris, C. R. et al. Array programming with NumPy. Nature 585, 357–362 (2020).

6. McKinney, W. Data Structures for Statistical Computing in Python. in 56–61 (Austin, Texas, 2010). doi:10.25080/Majora-92bf1922-00a.

7. Pedregosa, F. et al. Scikit-learn: Machine learning in python. J. Mach. Learn. Res. 12, 2825–2830 (2011).

8. Rideout, J. R. et al. biocore/scikit-bio: scikit-bio 0.5.9: Maintenance release. Zenodo 10.5281/ZENODO.8209901 (2023).

9. Hunter, J. D. Matplotlib: A 2D Graphics Environment. Comput. Sci. Eng. 9, 90–95 (2007).

10. Waskom, M. seaborn: statistical data visualization. J. Open Source Softw. 6, 3021 (2021).

11. Inc., P. T. Plotly python graphing library. (2015).

